# NBR1-mediated p62-liquid droplets enhance the Keap1-Nrf2 system

**DOI:** 10.1101/709105

**Authors:** Pablo Sánchez-Martín, Yu-shin Sou, Shun Kageyama, Masaaki Komatsu

**Author notes:** Correspondence should be addressed to Masaaki Komatsu.

## Abstract

p62/SQSTM1 is a multivalent protein that has an ability to cause a liquid-liquid phase separation and serves as a receptor protein that participates in cargo isolation during selective autophagy. This protein is also involved in the non-canonical activation of the Keap1-Nrf2 system, a major oxidative stress response pathway. Here we show a role of Neighbor of BRCA1 gene 1 (NBR1), an autophagy receptor structurally similar to p62/SQSTM1, in the p62-liquid droplet formation and the Keap1-Nrf2 pathway. The overexpression of NBR1 blocked selective degradation of p62/SQSTM1 through autophagy and promoted the accumulation and phosphorylation of p62/SQSTM1 in liquid-like bodies, which is required for the activation of Nrf2. NBR1 was induced in response to oxidative stress, and then the p62-mediated Nrf2 activation was up-regulated. Conversely, loss of *Nbr1* suppresses not only the formation of p62/SQSTM1-liquid droplets but also p62-dependent Nrf2 activation during oxidative stress. Taken together, our results show that NBR1 mediates p62/SQSTM1-liquid droplet formation to activate the Keap1-Nrf2 pathway.

## Introduction

When the concentration of molecules reaches a threshold, or when some modification increases the valency, liquid-liquid phase-separation occurs, and similar components gather and concentrate to be able to promote biochemical reactions or to sequester unwanted molecules. Such structures are called liquid droplets as they have liquid-like properties, and the inside is a reversible structure in which fluidity and biochemical reactions are kept (Dolgin, 2018). There are many droplets of this kind in the cell such as stress granules, P-granules and nucleoli, and they control the integrated stress response mechanism(s), that is, the robustness that protects cells from harmful disturbances like environmental stresses (Banani, Lee, Hyman, & Rosen, 2017). Liquid droplets are dynamic spherical structures that undergo fusion and fission events and are comparable in size to organelles. For this reason they are often referred to as membraneless organelles (Boeynaems et al., 2018). Excessive or abnormal droplets should be dispersed or be excluded adequately and promptly. Therefore, it is plausible that autophagy pathway mediates their turnover (Buchan, Kolaitis et al., 2013, Sun, Wu et al., 2018, Zhang, Yan et al., 2009). In fact, stress granules and P-granules are degraded by autophagy (Buchan, Kolaitis, Taylor, & Parker, 2013; Zhang et al., 2009). Meanwhile, suppression of autophagy is accompanied by the accumulation of structures in which liquid droplets are gelated and/or aggregated (Mizushima & Komatsu, 2011).

Macroautophagy (hereafter referred to as autophagy) is a cytoprotective mechanism that provides eukaryotic cells with building blocks for anabolic functions (Kaur & Debnath, 2015, Mizushima & Komatsu, 2011). In brief, after the appearance of a stressful stimulus (metabolic stress, organelle damage, and invasive microbes), a double membrane arises to isolate the cytosolic material in a compartment called the autophagosome. The autophagosome then fuses with a lysosome, promoting the degradation of the sequestered materials (Dikic & Elazar, 2018). While starvation-induced autophagy is thought to randomly degrade cytosolic components, under certain circumstances, autophagosomes selectively surround and degrade specific cargoes. The latter is called selective autophagy and contributes to the cellular homeostasis by degrading soluble proteins, liquid droplets, protein aggregates, excess or degenerated organelles and invasive microbes (Gatica, Lahiri et al. 2018). The molecular mechanism of autophagosome membrane formation in conventional and selective autophagy is considered to be common, but in selective autophagy, the “label of each cargo” or the presence of a “receptor protein” ensures the selectivity. “Labeling of each cargo” means ubiquitination of cargo or localization of receptor protein to cargo. “Receptor protein” refers to a group of proteins that bind to the cargo and to the autophagosomal proteins, the LC3 or GABARAP family. Receptor proteins are divided into ubiquitin-binding receptor proteins that recognize the ubiquitin chain of cargo, and cargo-localized receptor proteins that localize directly on the cargo. Both types of receptor proteins have LC3-interacting motif (LIR) or GABARAP-interacting motif (GIM) and bind directly to the LC3-, the GABARAP family, or both (Mizushima, 2018).

p62/SQSTM1 (hereafter referred to as p62) is a stress-inducible protein able to change among binding partners and cellular localizations, and it forms liquid-droplet structures in a context-dependent manner. This protein is mainly defined as a ubiquitin-binding receptor protein for selective autophagy (Sanchez-Martin & Komatsu, 2018). Besides this role, its ability to interact with multiple binding partners allows p62 to act as a main regulator of the activation of the Nrf2, mTORC1 and NF-κB signaling pathways, linking p62 to the oxidative defense system, nutrient-sensing and inflammation, respectively (Moscat, Karin et al., 2016, Sanchez-Martin, Saito et al., 2019). Recent reports revealed that once p62 binds to ubiquitin chains, it acquires liquid-like properties (Sun et al., 2018). Such phase-separated droplets allow the exchange of their components, including ubiquitin and LC3, with the surrounding environment. Consequently, the droplets can also function as nodes from which signaling cascades can be activated in the context of selective autophagy.

Here, we show a role of Neighbor of BRCA1 gene 1 (hereafter referred to as NBR1), which serves as a ubiquitin-binding autophagy receptor and is a binding partner of p62, in the p62-liquid droplets formation and the p62-mediated Nrf2 activation. Increased levels of NBR1 stabilized p62 and promoted the formation of liquid droplets resistant to autophagic degradation, leading to a robust induction of Nrf2 targets including p62. Meanwhile, loss of *NBR1* attenuated the p62-mediated activation of Nrf2 in response to oxidative stress. Our results indicate that the collaboration between p62 and NBR1 stretches beyond their role as selective autophagy receptors and that NBR1 is a new player in the orchestration of the antioxidant cell response.

## Results

### NBR1 increases the formation of p62-liquid droplets

Autophagy receptors p62 and NBR1 have been shown to act cooperatively in different forms of selective autophagy (Kirkin, Lamark et al., 2009a, Lamark, Kirkin et al., 2009). This cooperation is also observed between p62 and other cargo receptors (Cemma, Kim et al., 2011, Zhang, Varela et al., 2019). However, it is currently unknown if autophagy receptors have a direct effect on each other. As p62 and NBR1 share a common domain distribution and directly interacts with each other, we wondered if NBR1 influences the autophagic and non-autophagic functions of p62. With that aim, we exogenously expressed NBR1 in mouse primary hepatocytes by the adenovirus system. The overexpression of NBR1 increased not only total p62 protein, but also its Serine 349 and Serine 403 phosphorylated forms (Fig. 1A) that are involved in the Nrf2-activation and the ubiquitin-binding followed by phase-separation, respectively. The gene expression of *p62*, which is mainly regulated by the transcription factors Nrf2, NF-κB and MiT/TFE was induced upon the overexpression of NBR1 (Fig. 1B). Double-immunofluorescent analysis with anti-p62 and anti-NBR1 antibodies was employed as a complementary approach to confirm the interplay between NBR1 and p62. As previously described (Kirkin, Lamark et al., 2009b), both proteins extensively co-localized in same punctate structures at endogenous levels, and the expression of NBR1 increased the size and number of such structures (Fig. 1C). According to two recent reports (Sun et al., 2018, Zaffagnini, Savova et al., 2018), the p62-positive structures possess liquid-like properties that are essential for its role as an autophagy receptor. To check if the structures formed by NBR1 overexpression are also liquid-like structures, we measured the exchange of components between the structures and the surrounding environment by a fluorescence recovery after photobleaching (FRAP) assay. To do this, we overexpressed C-terminal GFP-tagged NBR1, NBR1-GFP in wild-type primary mouse hepatocytes. Similar to non-tagged NBR1, NBR1-GFP completely co-localized with p62-positive structures (Supplementary Fig. S1). The signal intensity of the structures positive for NBR1-GFP was recovered at 8-12 min after photobleaching (Fig. 1D and Supplementary Video S1), indicating that the p62- and NBR1-positive structures have liquid-like properties.

**Figure 1.**
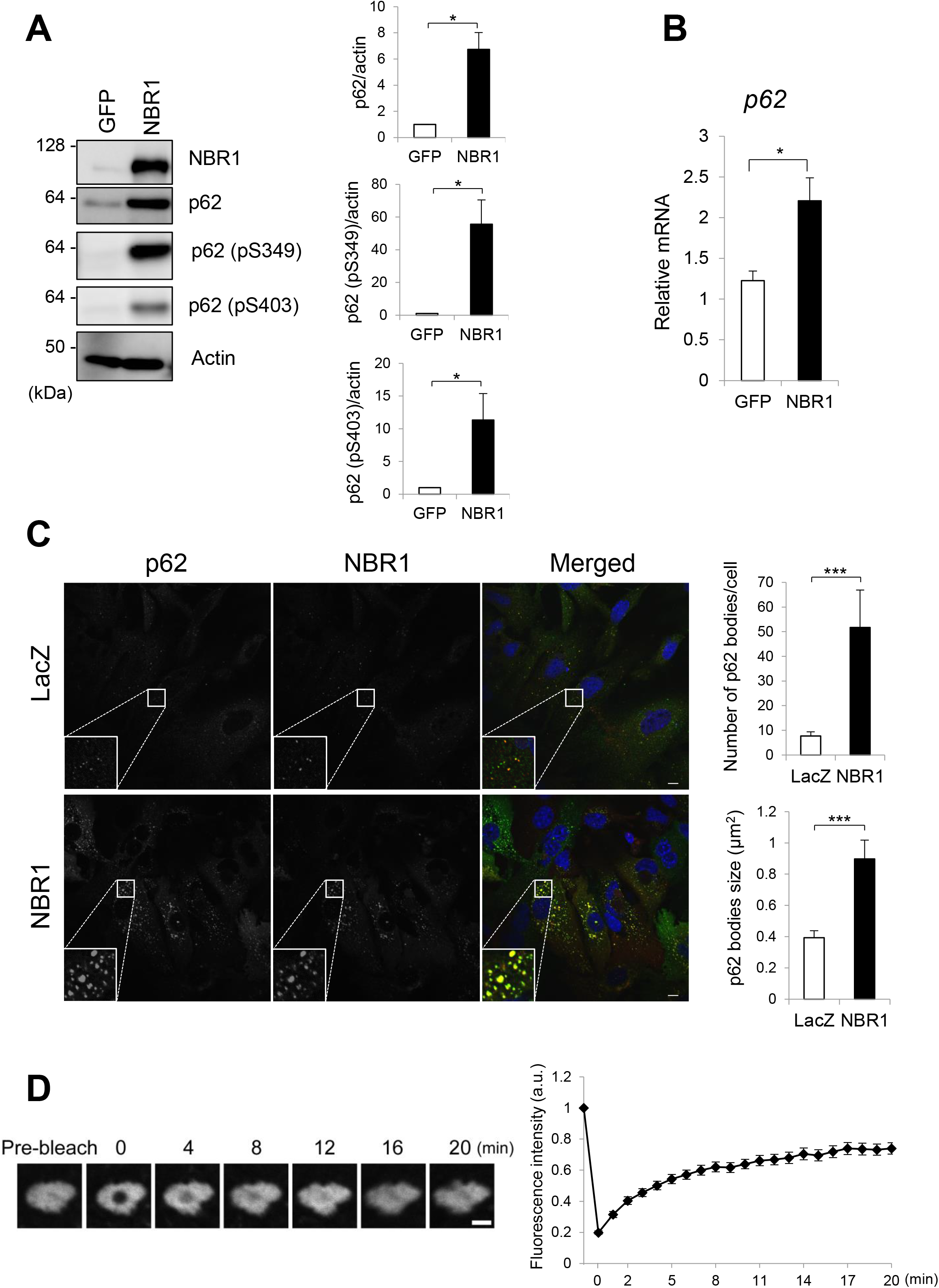
NBR1 promotes an increase in p62-levels, phosphorylation and body formation. (**A**) Immunoblot analysis. Primary hepatocytes prepared from wild-type mice were infected with adenovirus against GFP or NBR1 for 48 hr. Cell lysates were prepared and subjected to immunoblot analysis with the indicated antibodies. Data shown are representative of three separate experiments. Bar graphs indicate the quantitative densitometric analysis of the indicated proteins relative to actin. Data are means ± s.e. \**P* < 0.05 as determined by Welch’s *t*-test. (**B**) Real-time PCR. Total RNAs were prepared from primary hepatocytes described in (A). Values were normalized against the amount of mRNA in the hepatocytes expressing GFP. The experiments were performed three times. Data are means ± s.e. \**P* < 0.05 as determined by Welch’s *t*-test. (**C**) Immunofluorescence microscopy. Wild type hepatocytes were infected with LacZ or NBR1 adenovirus for 48 hr and then immunostained with anti-p62 and anti-NBR1 antibodies. Each inset is a magnified image. Bars: 10 μm. The number and size of p62 bodies were measured in more than 100 cells. Data are means ± s.e. \*\*\**P* < 0.001 as determined by Welch’s *t*-test. (**D**) FRAP assay. Wild type hepatocytes cultured in glass-bottom plates were infected with GFP-NBR1 adenovirus for 48 hr. The signal recovery after photobleaching was measured and quantified.

### NBR1 regulates the Nrf2-Keap1 pathway in a p62-dependent manner

Since p62 is involved in the regulation of multiple signaling pathways including mTORC1, p38 MAP kinase and Nrf2 (Sanchez-Martin et al., 2019), we examined if p62-phase separation due to overexpression of NBR1 affects such pathways. Expression of NBR1 in primary mouse hepatocytes did not have any effect on the conversion of LC3-I to LC3-II that is commonly used as a marker for autophagy fitness (KlionskyAbdelmohsen et al., 2016) (Supplementary Fig. S2). The phosphorylation of S6K, a substrate of mTORC1 was not affected by the expression of NBR1 (Supplementary Fig. S2). We did not observe any change in phosphorylation of p38 representing activation of p38, by NBR1-expression (Supplementary Fig. S2). Meanwhile, an increase level of NAD(P)H dehydrogenase, quinone 1 (Nqo1), an Nrf2-target was detected in the hepatocytes expressing NBR1 (Supplementary Fig. S2), suggesting that NBR1 could be involved in the activation of this signaling pathway.

To confirm the role of NBR1 in the activation of Nrf2, we checked if NBR1 expression affected the nuclear translocation of Nrf2 that is a key step in the activation of the pathway (Suzuki & Yamamoto, 2017). As shown in Figure 2A, when NBR1 was expressed into primary mouse hepatocytes isolated from *p62*^*f*/*f*^ mice, the nuclear level of Nrf2 significantly increased compared with that in the hepatocytes expressing GFP. Expectedly, gene expression of Nrf2-targets such as *Nqo1*, *Glutamate-cysteine ligase catalytic subunit* (*Gclc*) and *UDP-glucose dehydrogenase* (*Ugdh*) was markedly induced by the overexpression of NBR1, but not in the case of overexpression of GFP (Fig. 2B). We verified up-regulation of Nqo1, Ugdh and Gclc proteins in the hepatocytes expressing NBR1 (Fig. 2A), implying the functional activation of Nrf2.

**Figure 2.**
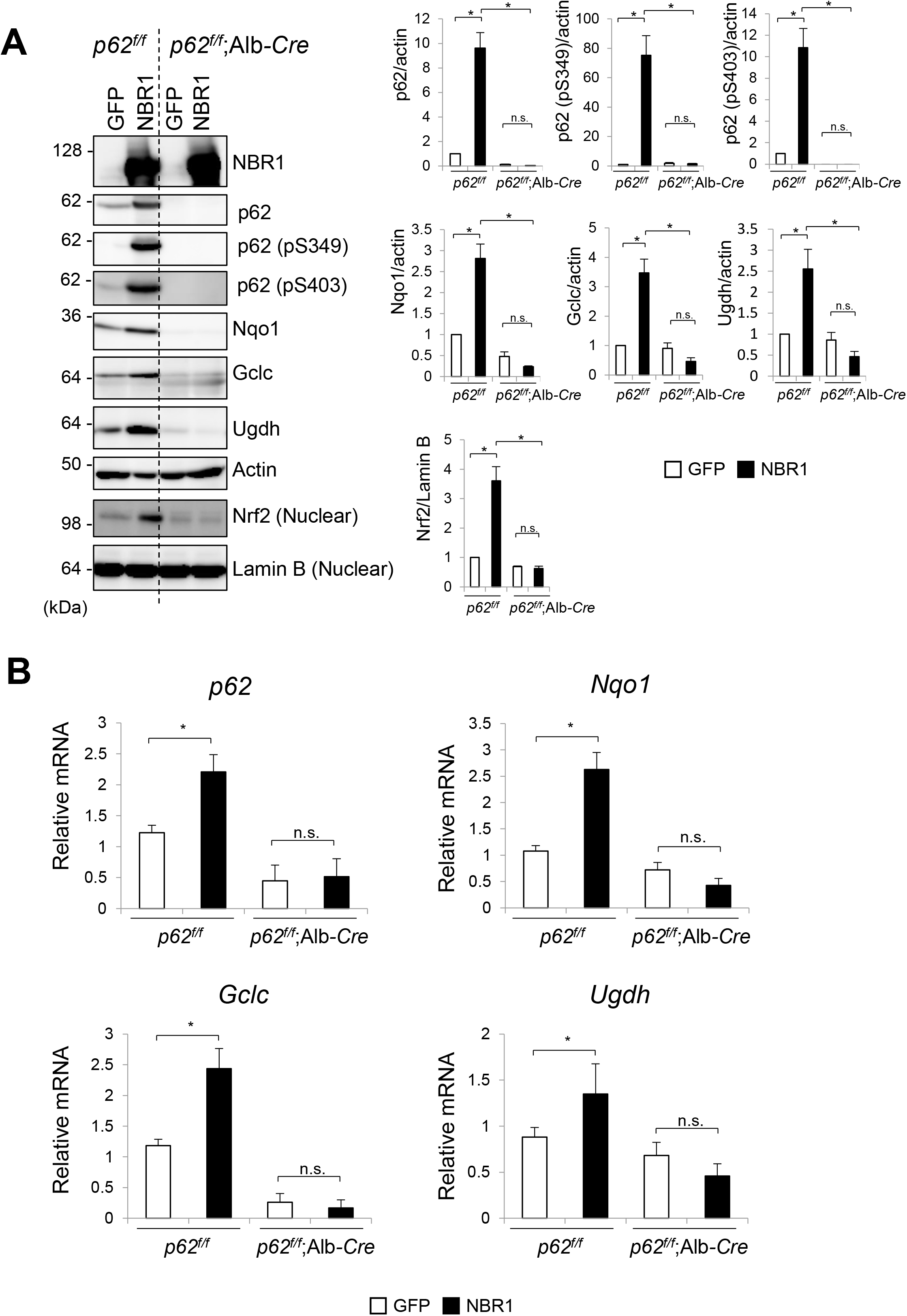
NBR1 increases the protein and mRNA levels of Nrf2 targets through p62. (**A**) Immunoblot analysis. Primary hepatocytes prepared from *p62*^*f*/*f*^ and *p62*^*f*/*f*^; Alb-*Cre* mice were infected with adenovirus against GFP or NBR1 for 48 hr. Cell lysates were prepared and subjected to immunoblot analysis with the indicated antibodies. Data shown are representative of three separate experiments. Bar graphs indicate the quantitative densitometric analysis of the indicated proteins relative to actin (cytosolic proteins) or Lamin B (nuclear proteins). Data are means ± s.e. \**P* < 0.05 as determined by Welch’s *t*-test. (**B**) Total RNAs were prepared from primary hepatocytes described in (A). Values were normalized against the amount of mRNA in the *p62*^*f*/*f*^ hepatocytes expressing GFP. The experiments were performed at least three times. Data are means ± s.e. \**P* < 0.05 as determined by Welch’s *t*-test.

p62 has the ability to activate Nrf2 under stress conditions (Komatsu, Kurokawa et al., 2010). To next examine the effect of p62 in the NBR1-mediated Nrf2 activation, we isolated hepatocytes from *p62*^*f*/*f*^ mice that expressed the Cre recombinase under the control of the Albumin promoter (*p62*^*f*/*f*^; Alb-*Cre*), resulting in the loss of the protein in the hepatocytes (Fig. 2A). Loss of p62 completely suppressed nuclear translocation of Nrf2 upon NBR1 expression (Fig. 2A). As a result, induction of Nrf2-target genes, which was observed in p62-competent hepatocytes was cancelled at both mRNA and protein levels by ablation of p62 (Fig. 2A and B). Collectively, these results indicate that NBR1 activates the Keap1-Nrf2 system in a p62-dependent fashion.

### NBR1 stabilizes the p62 protein by preventing its autophagic degradation

How does NBR1 achieve the accumulation of p62? Since *p62* is one of Nrf2-targets, we speculated that the NBR1-mediated Nrf2 activation contributes to this. To validate this idea, we isolated primary mouse hepatocytes from *Nrf2*^*f*/*f*^ and *Nrf2*^*f*/*f*^; Alb-*Cre* mice (Fig. 3A).

**Figure 3.**
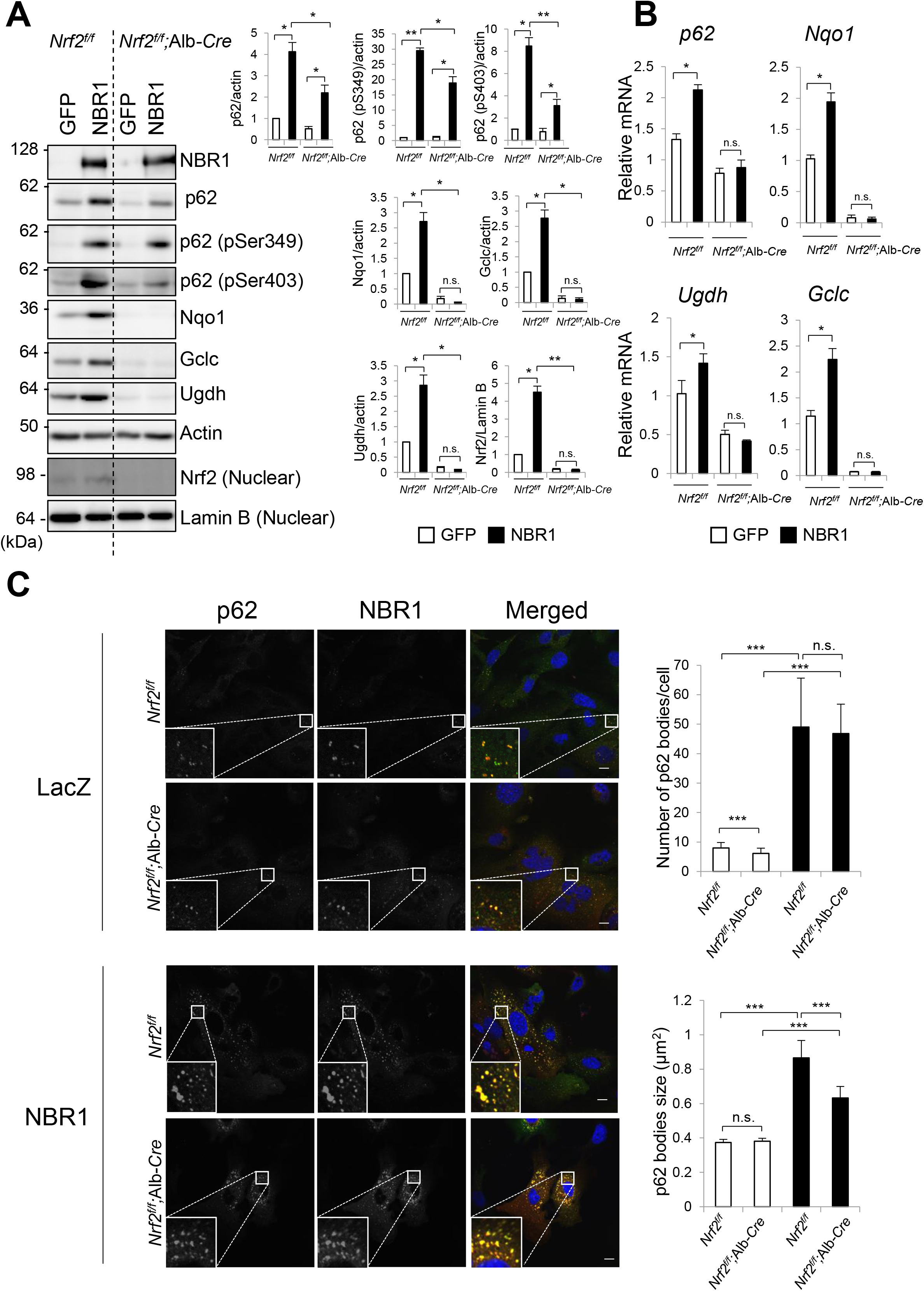
NBR1 enhances the protein levels of p62 partially independent on Nrf2. (**A**) Immunoblot analysis. Primary hepatocytes prepared from *Nrf2*^*f*/*f*^ and *Nrf2*^*f*/*f*^; Alb-*Cre* mice were infected with adenovirus against GFP or NBR1 for 48 hr. Cell lysates were prepared and subjected to immunoblot analysis with the indicated antibodies. Data shown are representative of three separate experiments. Bar graphs indicate the quantitative densitometric analysis of the indicated proteins relative to actin (cytosolic proteins) or Lamin B (nuclear proteins). Data are means ± s.e. \**P* < 0.05, and \*\**P* < 0.01 as determined by Welch’s *t*-test. (**B**) Real-time PCR. Total RNAs were prepared from primary hepatocytes described in (A). Values were normalized against the amount of mRNA in the *Nrf2*^*f*/*f*^ hepatocytes expressing GFP. The experiments were performed at least three times. Data are means ± s.e. \**P* < 0.05 as determined by Welch’s *t*-test. (**C**) Immunofluorescence microscopy. Hepatocytes isolated from *Nrf2*^*f*/*f*^ and *Nrf2*^*f*/*f*^; Alb-*Cre* mice were infected with LacZ or NBR1 adenovirus for 48 hr and then immunostained with anti-p62 and anti-NBR1 antibodies. Each inset is a magnified image. Bars: 10 μm. The number and size of p62 bodies were measured in more than 100 cells. Data are means ± s.e. \*\*\**P* < 0.001 as determined by Welch’s *t*-test.

Unexpectedly, the expression of NBR1 in hepatocytes lacking *Nrf2* still had an ability to increase the levels of both total p62 and its phosphorylated form albeit their levels were slightly but significantly lower than those in Nrf2-competent hepatocytes (Fig. 3A). Like gene expression of typical Nrf2-target genes, the induction of *p62*-transcript upon overexpression of NBR1 was repressed by loss of *Nrf2* (Fig. 3B). These results indicate that Nrf2 partially participates to the accumulation of p62 under NBR1-expressing conditions through induction of *p62*-transcript. Actually, we confirmed that NBR1 is able to up-regulate even the protein level of exogenously expressed p62 (Supplementary Fig. S3). The p62-liquid droplets were observed even in *Nrf2*-knockout hepatocytes, and the number and size increased by expression of NBR1 (Fig. 3C).

Although p62 is degraded through alternative pathways under certain situations (Mejlvang, Olsvik et al., 2018, Song, Li et al., 2016), this protein is mainly and selectively degraded by autophagy (Komatsu, Waguri et al., 2007). Thus, a possible explanation for the increase in the protein level of p62 observed in Figure 3A is that NBR1 prevents the autophagic degradation of p62. To examine this hypothesis, we conducted the autophagy-flux assay. The treatment of control GFP-expressing hepatocytes with Bafilomycin A_1_ (BafA_1_), an inhibitor of autophagosome-lysosome fusion caused increased levels of LC3-II, p62 and its phosphorylated forms (Fig. 4). As in control hepatocytes, we observed prominent accumulation of LC3-II in NBR1-expressing hepatocytes by the treatment of BafA_1_ (Fig. 4). In striking contrast to the control hepatocytes, the treatment of BafA_1_ did not further augment increased levels of p62 and its phosphorylated forms due to the NBR1-overexpression (Fig. 4). These results suggest that NBR1 inhibits selective autophagy of p62 rather than general autophagy.

**Figure 4.**
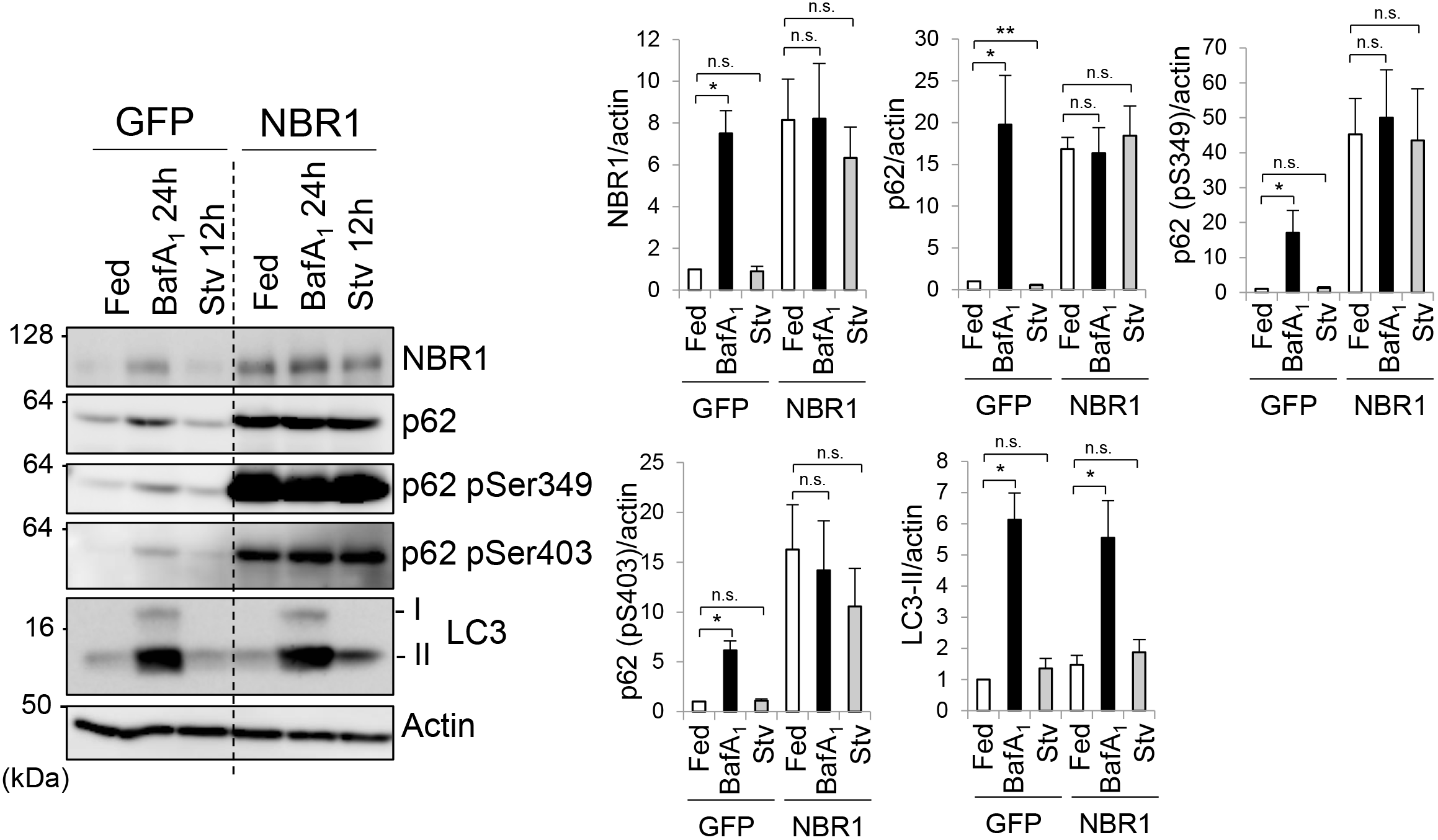
NBR1 prevents the autophagic degradation of p62. Immunoblot analysis. Primary hepatocytes prepared from wild-type mice were infected with adenovirus against GFP or NBR1 for 48 hr and subsequently treated with 100 nM Bafilomycin A_1_ (BafA_1_) for 24 hr or cultured in amino acid-deprived medium (Stv) for 12 hr. Cell lysates were prepared and subjected to immunoblot analysis with the indicated antibodies. Data shown are representative of three separate experiments. Bar graphs indicate the quantitative densitometric analysis of the indicated proteins relative to actin. Data are means ± s.e. \**P* < 0.05, and \*\**P* < 0.01 as determined by Welch’s *t*-test.

### NBR1 is a stress-inducible protein indispensable for full activation of the p62-Keap1-Nrf2 system

We have shown a role of NBR1 in the p62-mediated Nrf2 activation. While gene expression of *p62* is induced in response to different forms of stress (Jain, Lamark et al., 2010, Ling, Kang et al., 2012, Settembre, Di Malta et al., 2011), the expression pattern of *NBR1* largely remains unclear. To investigate the expression of endogenous NBR1 upon the exposure of oxidative stress, we selected the treatment of sodium arsenite (As [III]) as an oxidative stimulus since it is known to activate Nrf2 and induce the expression of *p62* (Shah, Trinh et al., 2017). The amount of NBR1 protein dramatically increased when primary mouse hepatocytes were exposed to As (III) (Fig. 5A). The increase in NBR1 was observed even in the absence of *p62* or *Nrf2* (Fig. 5A). Real-time PCR analysis revealed that Nrf2 is not responsible for gene expression upon the exposure of As (III) (Fig. 5B). This suggested that, despite its similarities, the induction of NBR1 during oxidative stress relies in a different transcription factor(s). In agreement with the case of NBR1-overexpression, the As (III)-mediated induction of NBR1 increased the size and number of p62-liquid droplets (Fig. 5C).

**Figure 5.**
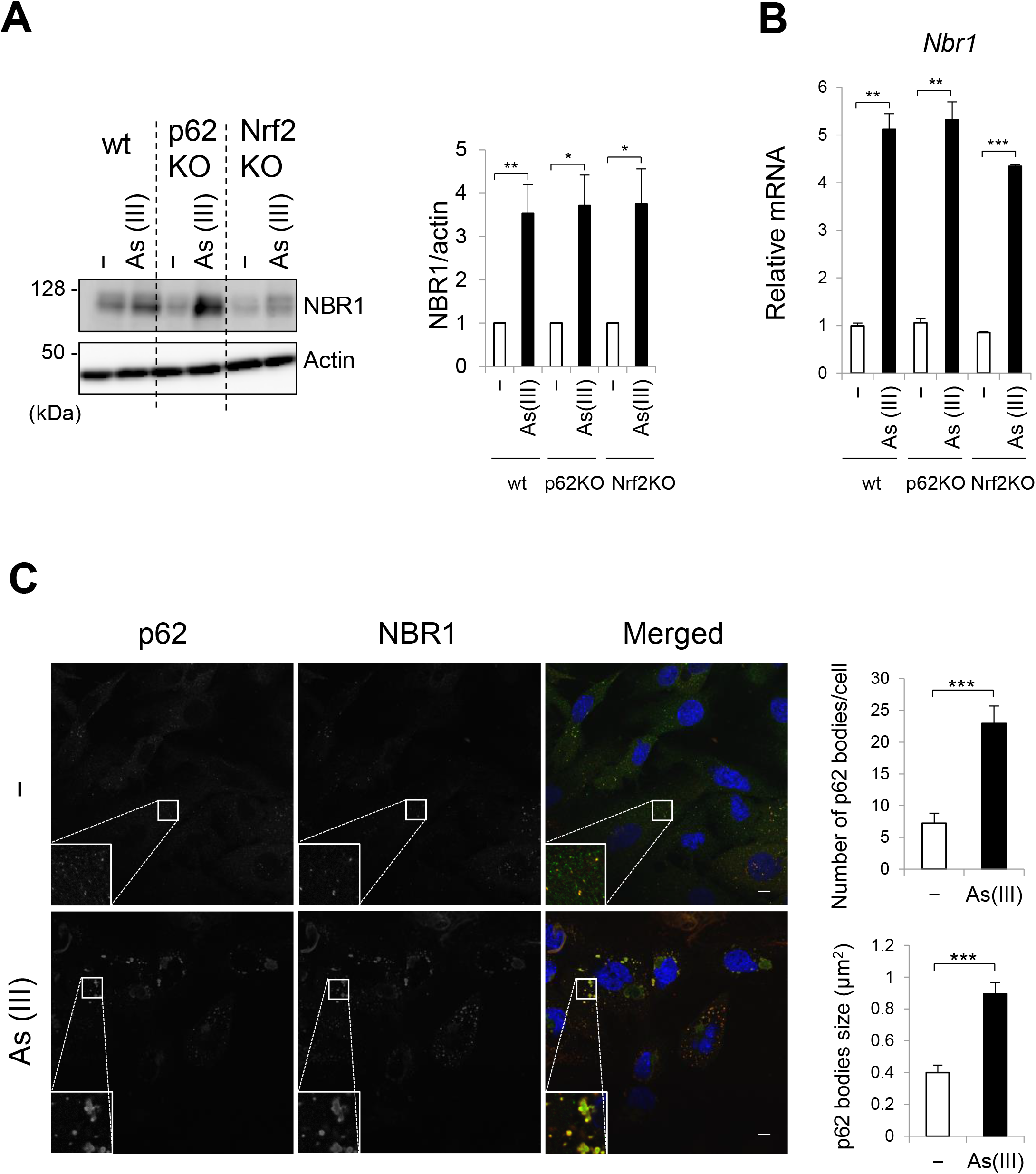
NBR1 is induced by oxidative stress. (**A**) Immunoblot analysis. Primary hepatocytes were isolated from wild-type (wt), *p62*^*f*/*f*^; Alb-*Cre* (p62KO) and *Nrf2*^*f*/*f*^; Alb-*Cre* (Nrf2KO) mice. The hepatocytes were challenged with 10 μM sodium arsenite [As (III)] for 12 hr. Cell lysates were prepared and subjected to immunoblot analysis with anti-NBR1 and anti-Actin antibodies. Data shown are representative of three separate experiments. Bar graphs indicate the quantitative densitometric analysis of the indicated proteins relative to actin. Data are means ± s.e. \**P* < 0.05, and \*\**P* < 0.01 as determined by Welch’s *t*-test. (**B**) Real-time PCR. Total RNAs were prepared from primary hepatocytes described in (A). Values were normalized against the amount of mRNA in each non-treated hepatocytes. (**C**) Immunofluorescence microscopy. Primary hepatocytes isolated from wild-type mice were challenged with 10 μM sodium arsenite [As (III)] for 12 hr and then immunostained with anti-p62 and anti-NBR1 antibodies. Each inset is a magnified image. Bars: 10 μm. The number and size of p62 bodies were measured in more than 100 cells. Data are means ± s.e. \*\*\**P* < 0.001 as determined by Welch’s *t*-test.

To assess the physiological relevance of NBR1 in the p62-dependent Nrf2 activation, we generated a line of conditional knockout mice for *Nbr1*, in which *Nbr1* is deleted in a Cre recombinase dependent manner (Supplementary Fig. S4A and B). Mice homozygous for the *Nbr1*^*flox*^ allele (referred to as *Nbr1*^*f*/*f*^ mice) were bred with a line of transgenic (Tg) mice, EIIa-*Cre* Tg that express the Cre recombinase under the control of the EIIa promoter in early embryo. The heterozygous mice (referred to *Nbr1*^+/−^) were obtained by crossbreeding of *Nbr1*^f/+^; EIIa-*Cre* with C57BL mice. We isolated mouse embryonic fibroblasts (MEFs) from *Nbr1*^+/+^, *Nbr1*^+/−^ and *Nbr1*^−/−^ mice and confirmed that neither *Nbr1-*transcript nor NBR1 protein was detectable in *Nbr1*^−/−^ MEFs (Supplementary Fig. S4C and D), indicating the effective recombination of the *Nbr1*^*flox*^ allele in the conditional knockout mice. Next, we generated hepatocyte-specific *Nbr1*-knockout mice, *Nbr1*^*f*/*f*^; Alb-*Cre* by crossbreeding *Nbr1*^*f*/*f*^ with an Alb-*Cre* Tg mice. Primary mouse hepatocytes were isolated from each wild-type, *p62*^*f*/*f*^; Alb-*Cre* and *Nbr1*^*f*/*f*^; Alb-*Cre* mice. Immunofluorescent analysis with anti-LC3 antibody showed appearance of LC3-positive dots corresponding to autophagosomes, and the number increased under nutrient-deprived conditions regardless genotypes (Supplementary Fig. S5A). We then examined the degradation of long-lived proteins. In wild-type hepatocytes, nutrient deprivation significantly enhanced the degradation, which was suppressed by the addition of lysosomal inhibitors, E64d and pepstatin A (EP) (Supplementary Fig. S5B). Degradation of long-lived proteins was also observed in *Nbr1*-deficient hepatocytes, and the level was comparable to with that in wild-type and *p62*-deficient hepatocytes (Supplementary Fig. S5B). These results indicate that loss of *Nbr1* does not affect general autophagy.

Finally, we sought whether stress-induced NBR1 expression contributes to the p62-mediated Nrf2 activation or not. With this aim, primary hepatocytes isolated from *Nbr1*^*f*/*f*^ mice were infected by adenoviral vector for Cre recombinase or GFP as a negative control. NBR1 in the hepatocytes was completely deleted by the expression of Cre recombinase, but not GFP (Fig. 6A). We observed the accumulation of total and phosphorylated p62 in both NBR1-competent and incompetent hepatocytes when challenged with As (III), but the induction levels in *Nbr1*-deficient hepatocytes were quite lower than control hepatocytes (Fig. 6A). More importantly, gene expression of Nrf2-targets upon the exposure of As (III) was significantly suppressed by loss of *Nbr1* (Fig. 6B). Furthermore, deletion of *Nbr1* resulted in a slight but significant reduction on the number and size of p62 liquid droplets during basal conditions, which was exacerbated when the hepatocytes were subjected to the treatment of As (III) (Fig. 6C). Taken together, these results indicate that NBR1 is an oxidative stress inducible protein and needed for full activation of the p62-Keap1-Nrf2 pathway.

**Figure 6.**
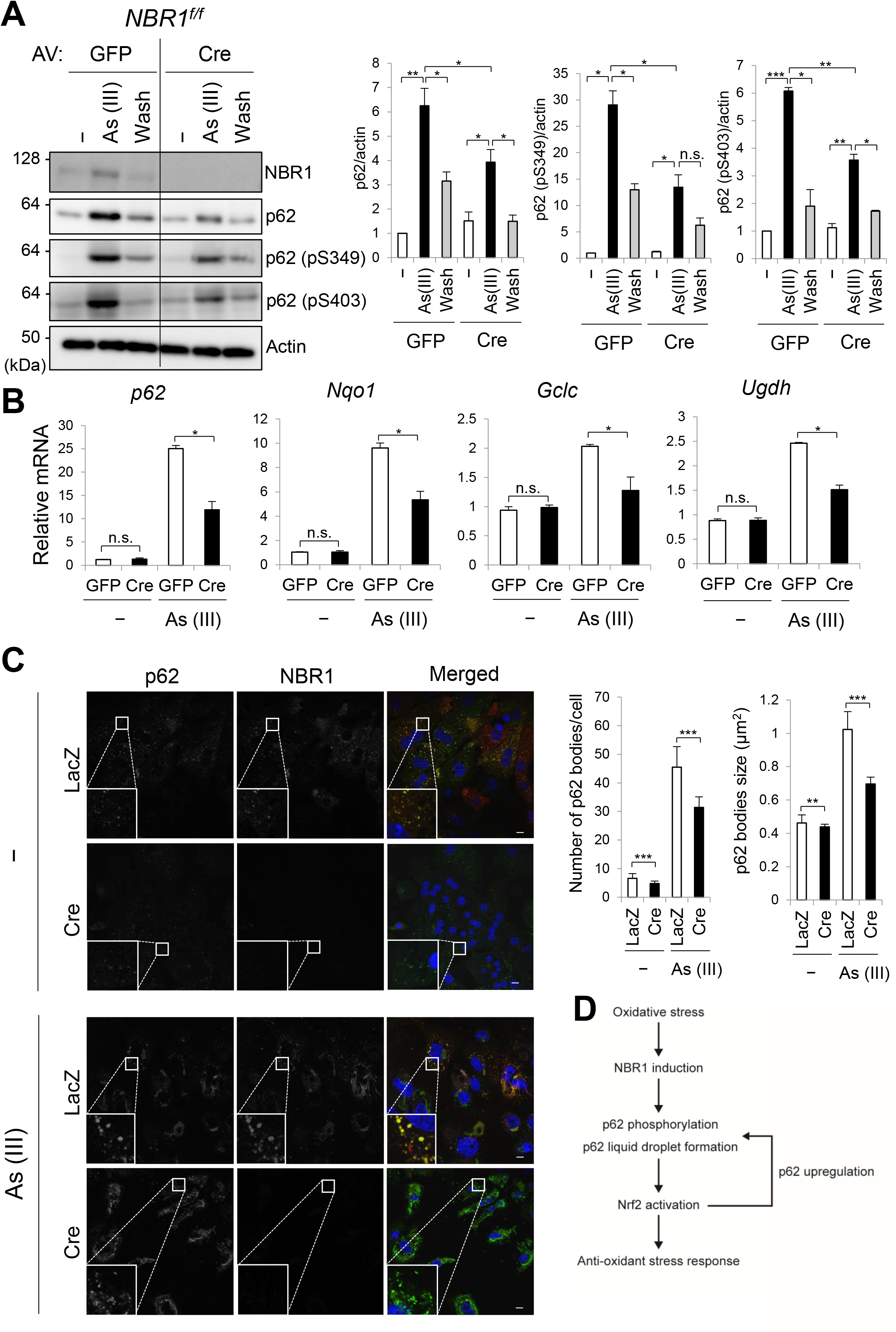
Deletion of NBR1 reduces the p62-mediated Nrf2 activation. (**A**) Immunoblot analysis. Primary hepatocytes were isolated from *Nbr1*^*f*/*f*^ mice and then infected with adenovirus GFP or Cre for 48 hr. Thereafter, the hepatocytes were divided into 3 groups: 1. Cultured in regular medium (−), 2. Treated with 10 μM As (III) for 12 hr [As (III)] and 3. After treated with 10 μM As (III) for 12 hr, cultured in regular medium for another 12 hr (Wash). Cell lysates were prepared and subjected to immunoblot analysis with indicated antibodies. Data shown are representative of three separate experiments. Bar graphs indicate the quantitative densitometric analysis of the indicated proteins relative to actin. Data are means ± s.e. \**P* < 0.05, \*\**P* < 0.01, and \*\*\**P* < 0.001 as determined by Welch’s *t*-test. (**B**) Real-time PCR. Total RNAs were prepared from primary hepatocytes described in (A). Values were normalized against the amount of mRNA in the *Nbr1*^*f*/*f*^ hepatocytes expressing GFP under normal culture conditions. The experiments were performed at least three times. Data are means ± s.e. \**P* < 0.05 as determined by Welch’s *t*-test. (**C**) Immunofluorescence microscopy. *Nbr1*^*f*/*f*^ hepatocytes were infected with LacZ or Cre adenovirus for 48 hr and, when indicated, challenged with 10 μM As (III) for 12 hr. The cells were then immunostained with anti-p62 and anti-NBR1 antibodies. Each inset is a magnified image. Bars: 10 μm. The number and size of p62 bodies were measured in more than 100 cells. Data are means ± s.e. \*\**P* < 0.01, and \*\*\**P* < 0.001 as determined by Welch’s *t*-test. (D) Role of NBR1 in the p62-liquid droplets and the p62-mediated Nrf2 activation. Oxidative stress up-regulates the level of NBR1, which enhances the phosphorylation and liquid-droplet formation of p62. The NBR1-containing p62-droplets are resistant to autophagic degradation and serve as signaling nodes for the activation of Nrf2. Since p62 is one of the targets of Nrf2, increased level of p62 in cooperation with NBR1 promote the formation of liquid droplets and contributes to the persistent activation of Nrf2 against oxidative stress.

## Discussion

A decade has passed since the identification of NBR1 as a selective autophagy receptor (Kirkin et al., 2009b), and yet little attention has been paid to its functional characterization. Previous research revealed that NBR1 and p62 interact with each other through their PB1 domains (Kirkin et al., 2009b) and act collaboratively in the autophagic clearance of protein aggregates (Kirkin et al., 2009b, Odagiri, Tanji et al., 2012), peroxisomes (Deosaran, Larsen et al., 2013) and invading pathogens (Shi, Fung et al., 2014). Herein we showed that NBR1 is induced in response to stress in order to promote the phase separation and phosphorylation of p62 and that the resulting droplets are resistant to autophagic degradation and serve as signaling hubs for the p62-mediated activation of Nrf2 (Fig. 6D).

In agreement with previous *in vitro* observations (Zaffagnini et al., 2018), NBR1 facilitated the formation of p62-liquid bodies in cells (Fig. 1). How does NBR1 promote the phase separation of p62? Since phase separation can happen when a concentration threshold of the components is reached, an easy explanation could be that the relative abundance of p62 and its binding partners surpass the concentration threshold, thus resulting in phase separation. As the overexpression of NBR1 increased the amounts of p62 via the activation of Nrf2 (Figs. 2 and 3), it is plausible that the concentration of components (*i.e.*, NBR1 and p62) is one factor influencing the formation of p62 bodies. Another possible mechanism is an increase in the valency of p62 by posttranslational modification. Posttranslational modifications including phosphorylation have been extensively identified as regulators of phase separation (Aumiller & Keating, 2016, Petri, Grimmler et al., 2007, Wippich, Bodenmiller et al., 2013). In matching with such criteria, the overexpression of NBR1 was accompanied by the robust phosphorylation of p62 at Serine 349 and Serine 403 (Fig. 1). Serine 349 phosphorylation is required for the competitive interaction with Keap1 (Ichimura, Waguri et al., 2013), while the phosphorylation of Serine 403 enhances the binding affinity of p62 to ubiquitin chains (Matsumoto, Wada et al., 2011), which is required for phase separation (Sun et al., 2018, Zaffagnini et al., 2018). Thus, it is likely that the increase in the phosphorylation states of p62 produced by NBR1 positively influence the formation of p62 bodies.

Remarkably, the p62 bodies induced by NBR1 overexpression were resistant to autophagic degradation (Fig. 4). Since the overexpression of NBR1 had no effect on the turnover of LC3-II, conventional autophagy is thought to be intact. How do these structures escape from autophagic degradation? The formation of the autophagosome along the interface of phase-separated p62 requires sequential and antagonistic steps. First, a complex between p62 and FIP200 is formed to initiate the selective autophagy of p62 (Turco, Witt et al., 2019). This complex is dependent on phosphorylation of p62 at Serine 349, which is located in close proximity to the LIR. This complex is mutually exclusive to the one formed between p62 and LC3, so the binding of LC3 occurs after dissociation from FIP200. Which step(s) does NBR1 inhibit? Since the overexpression of NBR1 enhanced both phase-separation of p62 and the phosphorylation of Serine 349 (Fig. 1), it is plausible that the p62-droplets are under priming for recruitment of FIP200. Thus, NBR1 might prevent the translocation of FIP200 onto the p62-bodies. Alternatively, the persistent phosphorylation of p62 at Serine 349 by the NBR1-overexpression may inhibit the binding transition from FIP200 to LC3-II. Actually, p62 is de-phosphorylated at Serine 349 in response to removal of As(III) (Eino, Kageyama et al., 2015), and the phospho-defective mutant of p62 is resistant to autophagy (Turco et al., 2019), suggesting the involvement of the de-phosphorylation in LC3-binding. Further analysis is needed to clarify how NBR1 regulates the removal of p62-liquid droplets.

What is the physiological role of the NBR1-mediated p62-bodies? We speculate that NBR1 shifts p62 bodies from autophagic cargo clusters to signaling nodes. Specifically, these signaling nodes seem specialized on the activation of the Keap1-Nrf2 antioxidant pathway, as other p62-related signaling pathways were hardly affected by the overexpression of NBR1 (Fig. S2). The fact that the NBR1-mediated activation of Nrf2 is lost in *p62*-deficient cells (Fig. 2) points out that the role of NBR1 in this pathway is achieved by increasing p62 protein levels and phosphorylation state. Most importantly, NBR1 was induced during oxidative stress in an Nrf2-independent manner (Fig. 5), implying that a rate-limiting step of NBR1-mediated Nrf2 activation is an induction of the expression of *NBR1*. *NBR1* is located on a region of chromosome 17q21.1 that is in close proximity to the *BRCA1* tumor suppressor gene. In fact, in mice, *Nbr1* and *Brca1* are separated by just less than 300 base pairs and share a common bidirectional promoter (Whitehouse, Chambers et al., 2004, Xu, Chambers et al., 1997). Besides an established role as tumor suppressor (Silver & Livingston, 2012), BRCA1 also participates in the orchestration of the oxidative stress response in multiple manners (Yi, Kang et al., 2014), including the stabilization of Nrf2 (Gorrini, Baniasadi et al., 2013). Potential transcription factors of the BRCA1/NBR1 bidirectional promoter include Sp1, C/EBPs and CREB (Xu et al., 1997), all of them involved in the antioxidant defense pathway (Lee, Cao et al., 2009, Ryu, Lee et al., 2003, Thon, Al Abdallah et al., 2010). Although further research is required to clarify the mechanism, one could envision that during oxidative threat, stress-inducible transcription factors would be recruited to the bidirectional promoter to secure the activation of the master antioxidant regulator Nrf2 by the action of both NBR1 and BRCA1.

## Material and Methods

### Mice

p62(Harada, Warabi et al., 2013) and Nrf2(Xue, Hou et al., 2013)-conditional knockout mice with C57BL/6 genetic background were previously described. Generation of Nbr1-conditional knockout mice was performed as in (Komatsu, Waguri et al., 2005). Mice were housed in specific pathogen-free facilities, and the Ethics Review Committee for Animal Experimentation of Niigata University and Juntendo University approved the experimental protocol.

### Isolation of mouse primary hepatocytes

Six-month-old mice were anesthetized with isoflurane, and the mouse livers were perfused by reserve flow through the vena cava using 20 mL of ex-perfusate solution (pH 7.4, 140 mM NaCl, 5mM KCl, 0.5 mM NaH_2_PO_4_, 10 mM HEPES, 4mM NaHCO_3_, 50 mM glucose and 0.5 mM GEDTA) and 30 mL of perfusate solution (pH 7.6, 70 mM NaCl, 6.7 mM KCl, 100 mM HEPES, 5 mM CaCl_2_, 35 units collagenase). After dissection, hepatocytes suspension was washed three times in HBSS (Gibco, Thermo Fisher Scientific) and the cells were grown in collagen-coated plates with William’s Medium E (Gibco, Thermo Fisher Scientific) supplemented with 10% fetal bovine serum (FBS).

### Immunoblot analysis

Cells were lysed in ice-cold TNE buffer (50 mM Tris-HCl, pH 7.5, 150 mM NaCl, 1 mM EDTA) containing 1% Triton X-100 and protease inhibitors. Nuclear and cytoplasmic fractions were prepared using the NE-PER Nuclear and Cytoplasmic Extraction Reagents (Thermo Fisher Scientific). Samples were separated by SDS-PAGE and then transferred to polyvinylidene difluoride (PVDF) membranes. Antibodies against NBR1 (4BR; Santa Cruz Biotechnology), p62 (GP62-C, Progen Biotechnik GmbH), LC3B (#2775, Cell Signaling Technology), T421/S424-phosphorylated S6K (#9204, Cell Signaling Technology), T389-phosphorylated S6K (#9205, Cell Signaling Technology), S240/S244-phosphorylated S6 (#2215, Cell Signaling Technology) T180/Y182-phosphorylated p38 (#9211, Cell Signaling Technology), Actin (MAB1501R, Merck Millipore Headquarters), Lamin B (M-20, Santa Cruz Biotechnology), Nqo1 (ab34173; Abcam), Ugdh (ab155005; Abcam), Gclc (ab41463, Abcam), Nrf2 (H-300; Santa Cruz Biotechnology), Keap1 (10503-2-AP; Proteintech Group) were purchased from the indicated suppliers and employed at 1:500 dilution. Anti-S349-phosphorylated p62 polyclonal antibody was raised in rabbits by using the peptide Cys+KEVDP(pS)TGELQSL as an antigen (Ichimura et al., 2013). Blots were then incubated with horseradish peroxidase-conjugated secondary antibody (Goat Anti-Mouse IgG (H + L), 115-035-166, Jackson ImmunoResearch; Goat Anti-Rabbit IgG (H + L) 111-035-144; Goat Anti-Guinea Pig IgG (H + L)) and visualized by chemiluminescence.

### Quantitative real-time PCR (qRT-PCR)

Using the Transcriptor First-Strand cDNA Synthesis Kit (Roche Applied Science, Indianapolis, IN, USA), cDNA was synthesized from 1 μg of total RNA. Quantitative PCR was performed using the LightCycler® 480 Probes Master mix (Roche Applied Science) on a LightCycler® 480 (Roche Applied Science). Signals were normalized against *Gusb* (β-glucuronidase). The sequences of the primers employed are provided in Supplementary Table 1.

### Immunofluorescence microscopy

Cells grown on coverslips were fixed in 4% paraformaldehyde in PBS for 10 min, permeabilized with 0.1% Triton X-100 in PBS for 5 min, blocked with 0.1% (w/v) gelatin (Sigma-Aldrich) in PBS for 45 min, and then incubated overnight with primary antibodies diluted 1:200 in gelatin/PBS. After washing, cells were incubated with Goat anti-Guinea pig IgG (H + L) Cross-Adsorbed Secondary Antibody, Alexa Fluor 488 (A11073, Thermo Fisher Scientific), and Goat anti-Mouse IgG (H + L) Highly Cross-Adsorbed Secondary Antibody, Alexa Fluor 647 (A21236, Thermo Fisher Scientific) at a dilution ratio of 1:1000 for 60 min. Cells were imaged using a confocal laser-scanning microscope (Olympus, FV1000) with a UPlanSApo ×60 NA 1.40 oil objective lens. The number and size of p62 bodies was analyzed with the ImageJ 1.51 software (Wayne Rasband, National Institutes of Health, USA).

To measure the fluorescence recovery after photobleaching (FRAP), cells were grown in 35mm glass base dishes (Iwaki, Japan). p62 bodies were bleached for 3 sec using a laser intensity of 70% at 480nm, and then the recovery was recorded for the indicated time.

### Protein degradation assay

The assay was performed essentially as described in (Komatsu et al., 2005). In brief, hepatocytes isolated from the indicated genotypes were plated at 5×10^4^ cells/well in collagen-coated 24-well plates and cultured in Williams’ E medium with 10% FBS for 24 hr. Cells were incubated with this medium also containing 0.5 μCi/ml [^14^C]leucine for 24 hr to label long-lived proteins. After washing with Williams’ E containing 2 mM of unlabeled leucine, cells were incubated with the medium for 2 hr to allow degradation of short-lived proteins and minimize the incorporation of labeled leucine. Next, the cells were washed again with PBS and incubated at 37°C with Krebs-Ringer bicarbonate medium and Williams’ E/10% FCS in the presence or absence of protease inhibitors (10 μg/ml E64d and pepstatin). After 4 hr, aliquots of the medium were taken and a one-tenth volume of 100% trichloroacetic acid was added to each aliquot. The mixtures were centrifuged at 12,000 g for 5 min, and the acid-soluble radioactivity was determined using a liquid scintillation counter. At the end of the experiment, the cultures were washed twice with PBS, and 1 ml of cold trichloroacetic acid was added to fix the cell proteins. The fixed cell monolayers were washed with trichloroacetic acid and dissolved in 1 ml of 1 N NaOH at 37°C. Radioactivity in an aliquot of 1 N NaOH was determined by liquid scintillation counting. The percentage of protein degradation was calculated according to the procedures published in (Gronostajski & Pardee, 1984).

## Acknowledgements

P.S.-M. is supported by a Grant-in-Aid for JSPS Research Fellows (P18099). Y.-S.S. is supported by a Grant-in-Aid for Scientific Research (C) (19K15043). S.K. is supported by a Grant-in-Aid for Early-Career Scientists (18K15043). M.K. is supported by a Grant-in-Aid for Scientific Research on Innovative Areas (19H05706), a Grant-in-Aid for Scientific Research (B) (18H02611), the Japan Society for the Promotion of Science (an A3 foresight program), and the Takeda Science Foundation (to M.K.).

## Conflict of Interests

We declare that we have no competing financial interests.

---

**Supplementary Figure S1.**
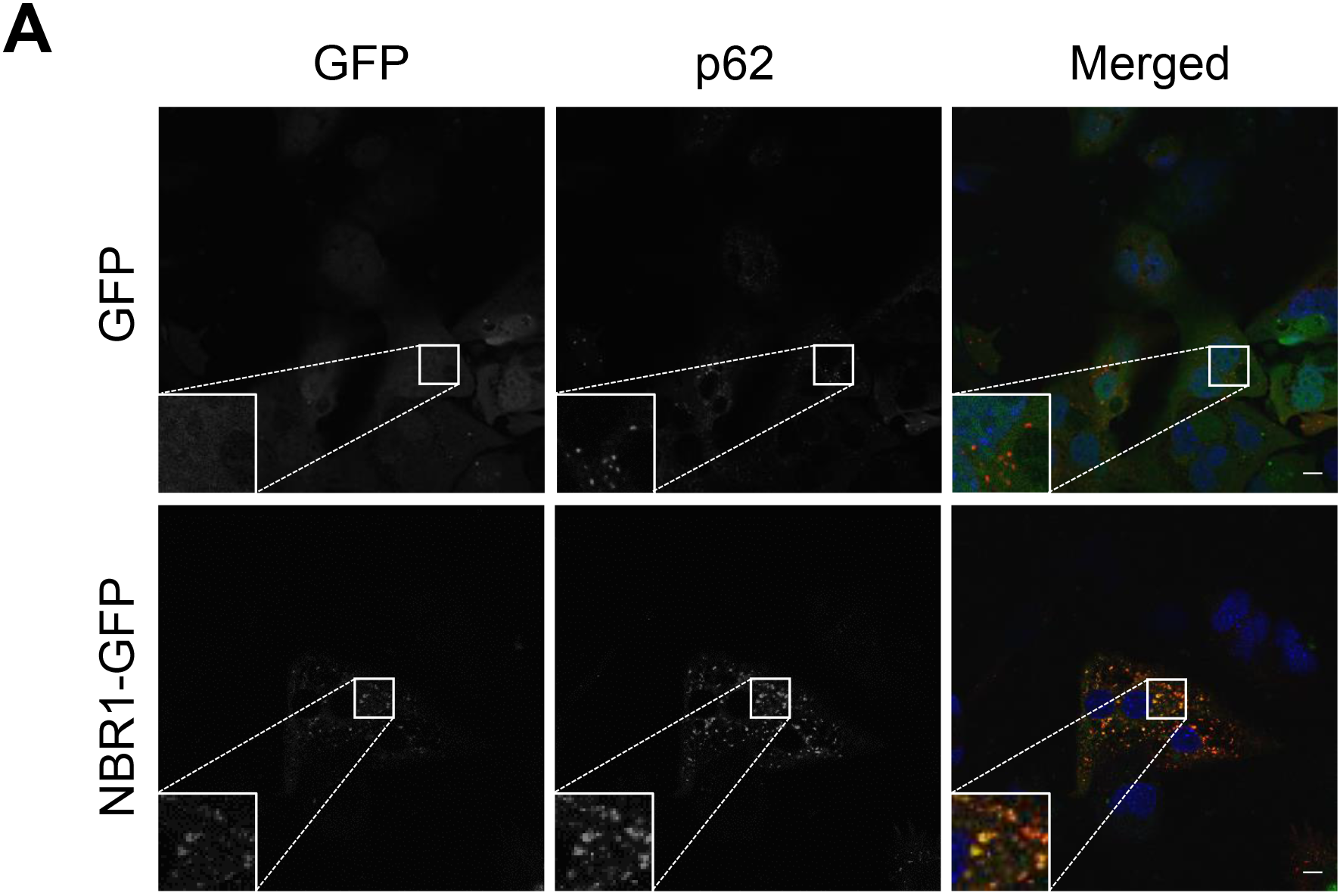
NBR1-GFP colocalizes with p62 bodies. Wild type hepatocytes were infected with GFP or NBR1-GFP adenovirus for 48 hr and then immunostained with anti-p62 antibody. Each inset is a magnified image. Bars: 10 μm.

**Supplementary Figure S2.**
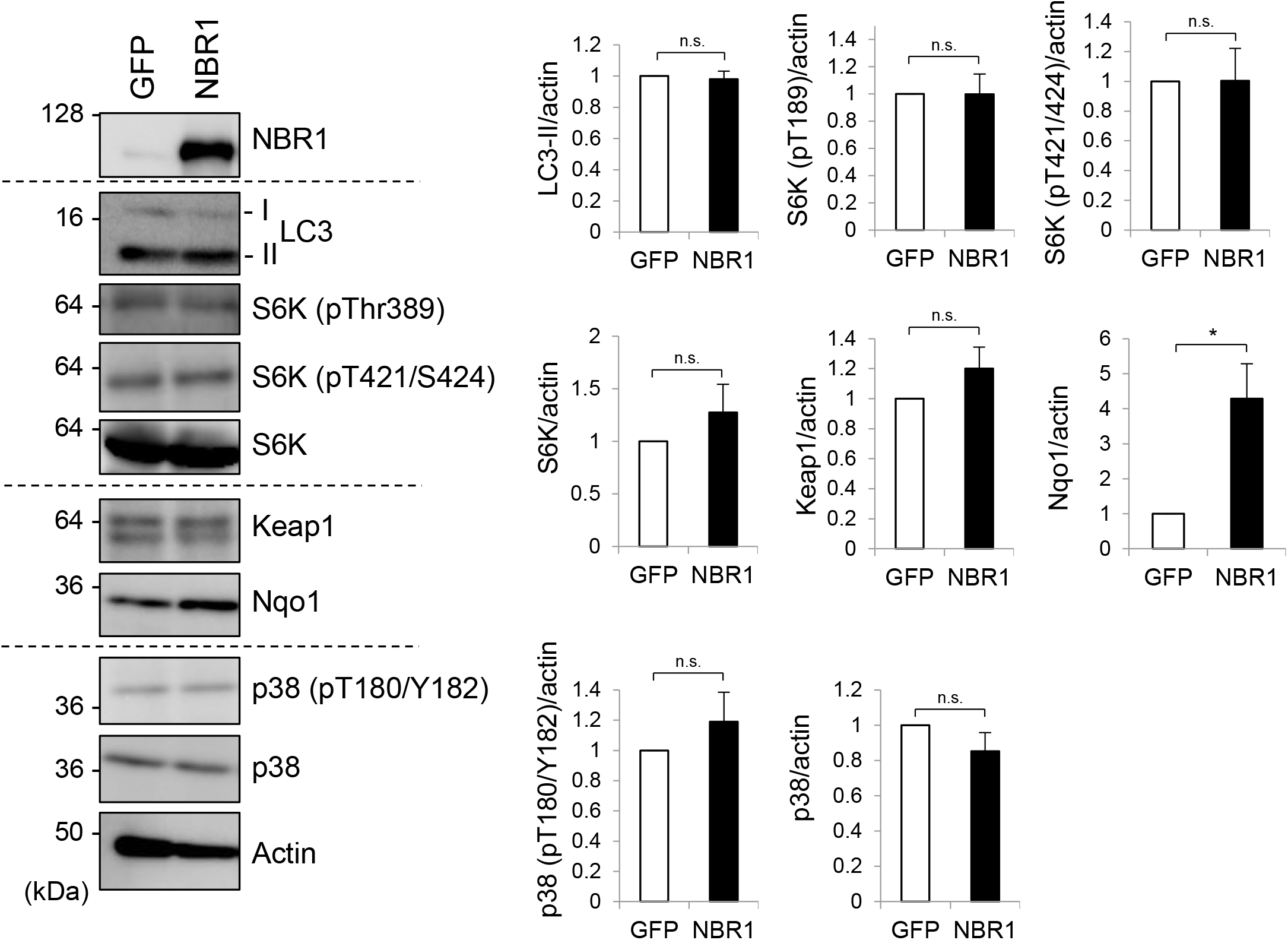
NBR1 influences Nrf2 activation but not other p62-related pathways. Primary hepatocytes prepared from wild-type mice were infected with adenovirus against GFP or NBR1 for 48 hr. Cell lysates were prepared and subjected to immunoblot analysis with the indicated antibodies. Data shown are representative of three separate experiments. Bar graphs indicate the quantitative densitometric analysis of the indicated proteins relative to actin. Data are means ± s.e. \**P* < 0.05 as determined by Welch’s *t*-test.

**Supplementary Figure S3.**
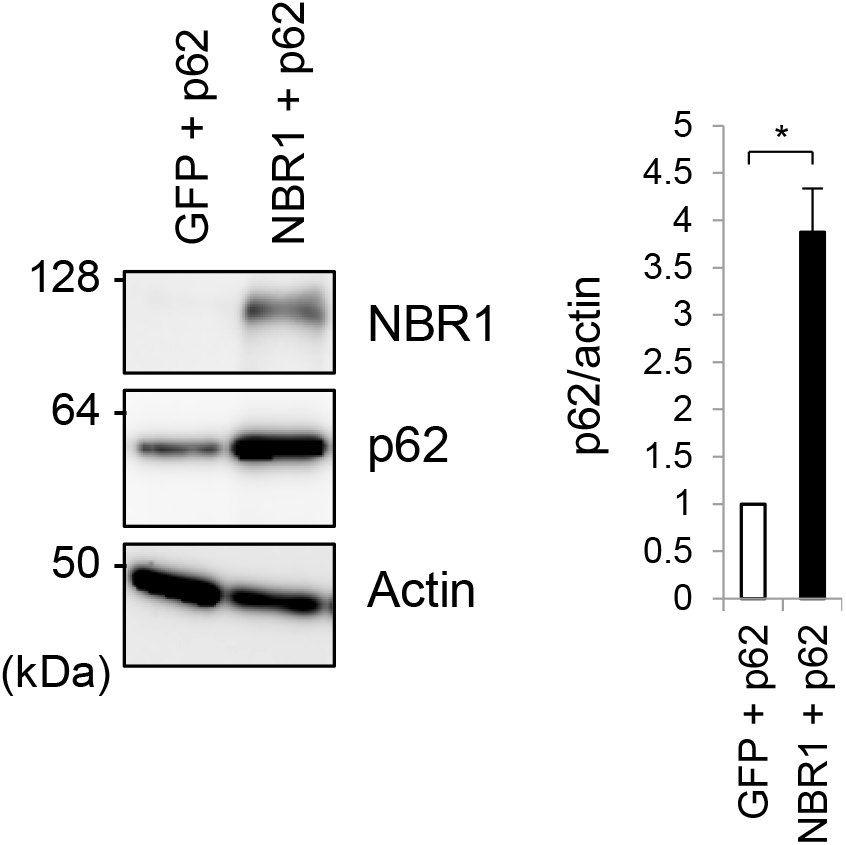
Overexpression of NBR1 promotes the accumulation of exogenous p62. Primary hepatocytes prepared from *p62*^*f*/*f*^; Alb-*Cre* mice were infected with adenovirus against GFP or NBR1 in combination with p62 for 48 hr. Cell lysates were prepared and subjected to immunoblot analysis with the indicated antibodies. Data shown are representative of three separate experiments. Bar graphs indicate the quantitative densitometric analysis of p62 relative to actin. Data are means ± s.e. \**P* < 0.05 as determined by Welch’s *t*-test.

**Supplementary Figure S4.**
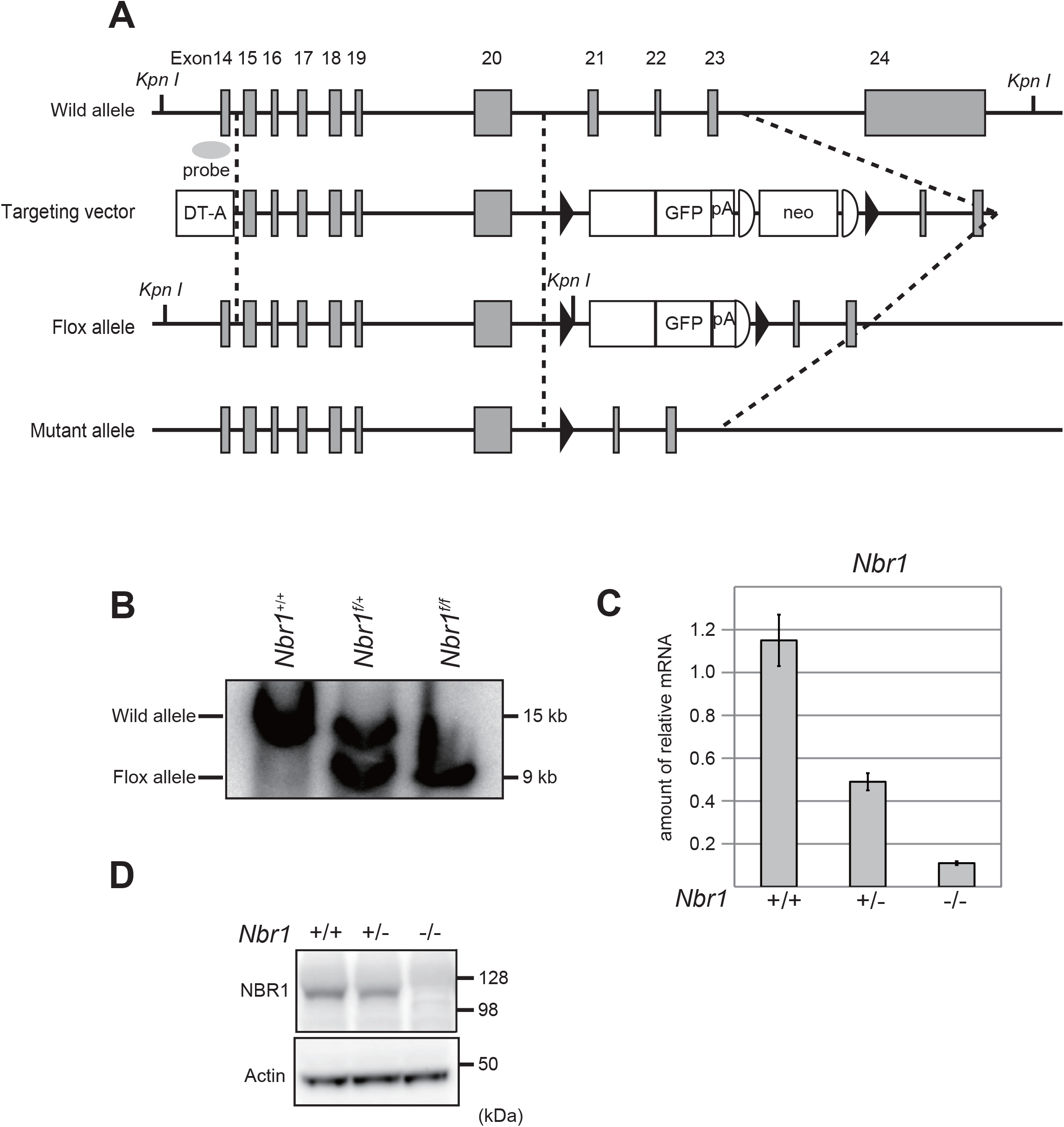
Generation of *Nbr1*-conditional knockout mice. (**A**) Schematic representation of the targeting vector and the targeted allele of the *Nbr1* gene. The coding exons numbered in accordance with the initiation site as exon 1 are depicted by grey boxes. Exon 21 was fused to a cDNA fragment encoded by exons 22, 23 and 24 (aa 2523-2964), GFP and polyA signal sequence was added. Neo resistant gene cassette (neo) with FLP sequences (representing semi-ellipsoid) was ligated behind the polyA signal sequence. The black triangles denote loxP sequence. A probe for Southern blot analysis is shown as a gray ellipse. (**B**) Southern blot analysis of genomic DNA extracted from mice tails. Wild-type and Flox alleles are indicated. (**C**) Expression of the *Nbr1* transcript in mouse embryonic fibroblasts (MEFs). Transcripts from the indicated genotypes were detected by real-time PCR analysis. (**D**) MEF cells from the indicated genotypes were subjected to immunoblot analysis with the anti-NBR1 and Actin antibodies. Data shown are representative of three separate experiments.

**Supplementary Figure S5.**
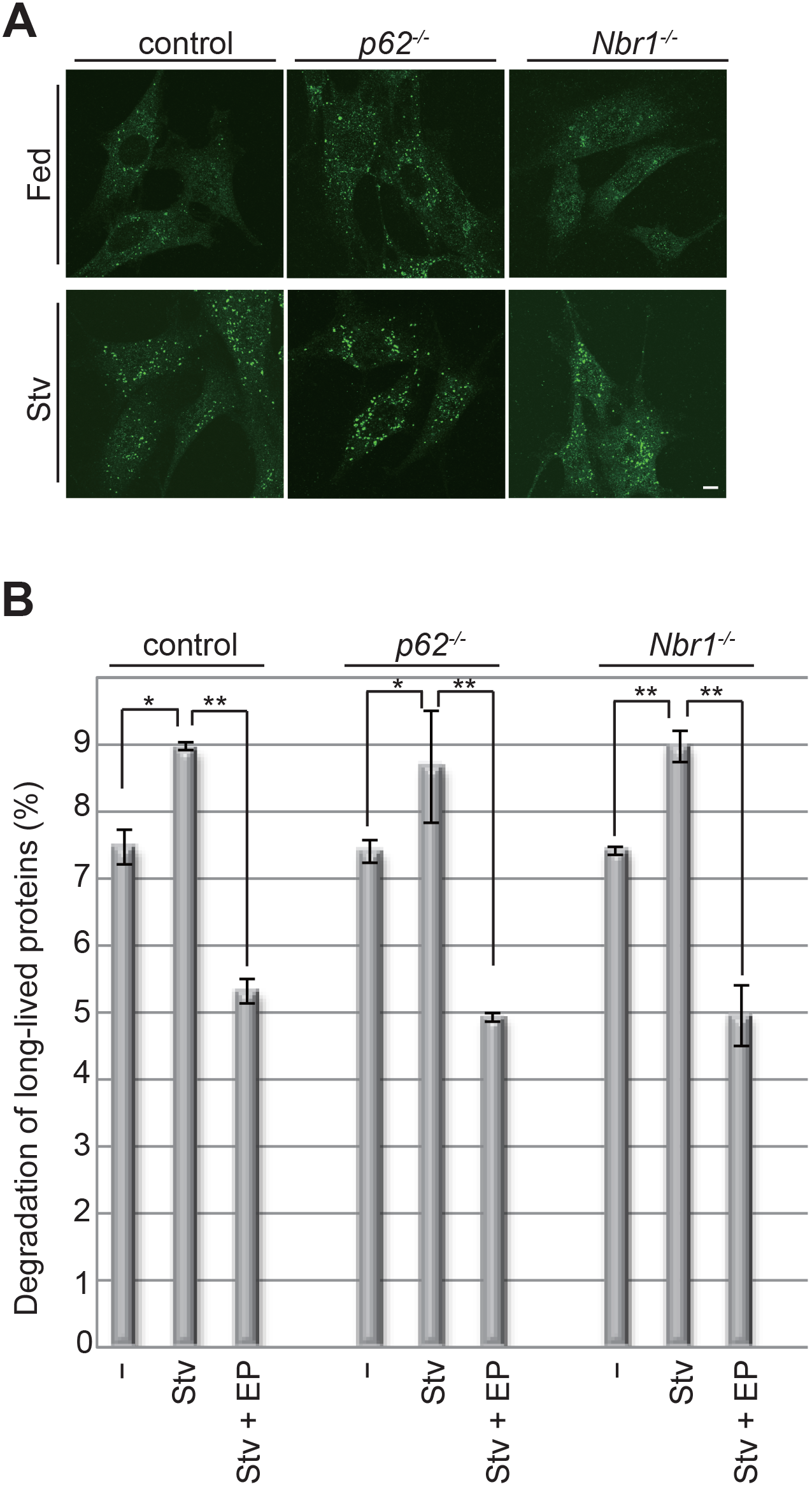
Autophagic activity in *Nbr1*-deficient hepatocytes. (**A**) Immunofluorescence microscopy. Primary hepatocytes from the indicated genotypes were isolated and subjected to immunostaining with an anti-LC3 antibody. Bars: 10 μm. (**B**) Hepatocytes from wild-type, *p62*- and *Nbr1*-knockout mice were isolated and labeled with [^14^C] leucine for 24 hr, and degradation of long-lived protein in deprived (Stv) or non-deprived (−) condition was measured. E64d and pepstatin (EP) was added as indicated. Data are means ± s.e. \**P* < 0.05, \*\**P* < 0.01 as determined by Welch’s *t*-test.

**Supplementary Video V1. NBR1-containig p62 structures have liquid-like properties.** Time lapse imaging of the signal recovery after photobleaching. Wild type hepatocytes cultured in glass-bottom plates were infected with GFP-NBR1 adenovirus for 48 hr.

**Supplementary Table 1.**
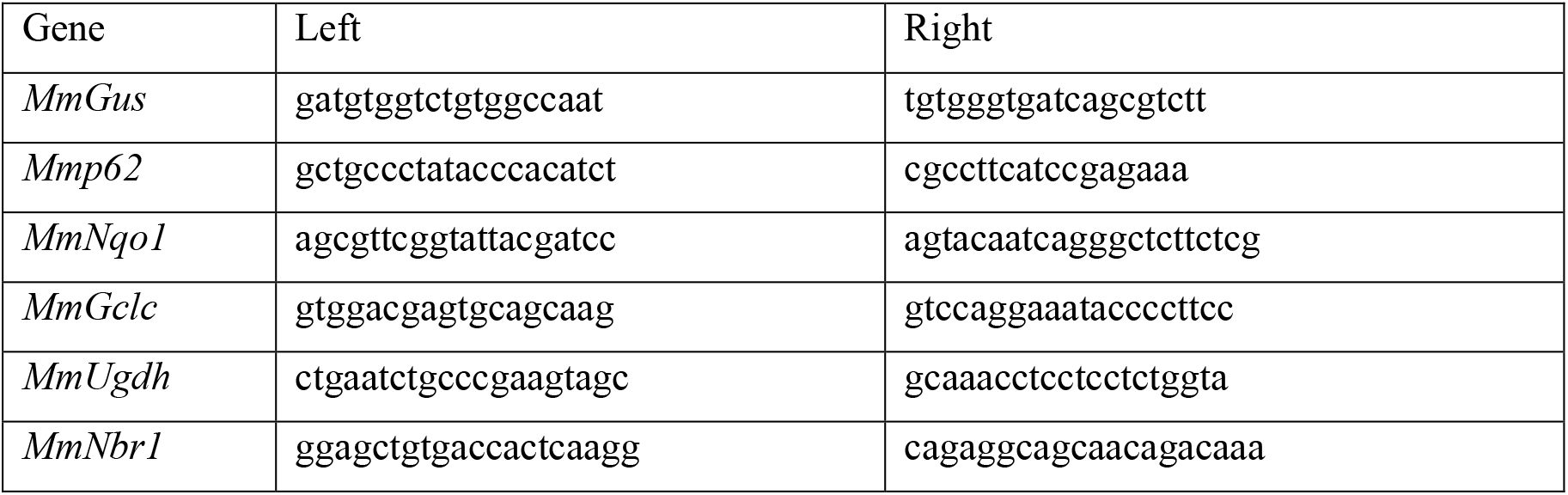
Primer sequences used in quantitative real-time PCR.

